# A degradable nanofibrous scaffold of poly(ε-Caprolactone-co-Lactide) for annulus fibrosus repair

**DOI:** 10.64898/2025.11.30.691409

**Authors:** Mansoor Chaaban, Chloé Falcoz, Cyrille Decante, Coline Pinese, Floriane Etienne, Paul Humbert, Maeva Dutilleul, Johann Clouet, Benjamin Nottelet, Christian Liebsch, Ann-Kathrin Greiner-Perth, Morten Vogt, Hans-Joachim Wilke, Xavier Garric, Jérôme Guicheux, Marion Fusellier, Catherine Le Visage

## Abstract

The intervertebral disc (IVD) is a key contributor to the spine’s biomechanical functions. It consists of a gelatinous core (nucleus pulposus, NP) surrounded by a fibrous ring (annulus fibrosus, AF). Accumulation of microcracks and tears in the AF can lead to NP herniation outside the disc space, compressing the nerve roots and causing pain. Herniation is a leading cause of low back pain and represents a major socioeconomic burden, with no effective regenerative treatment currently available. Previously, we demonstrated the potential of a poly(ε-caprolactone) (PCL)-based implant for the closure of an annulus fibrosus (AF) defect. However, the slow in vivo degradation of PCL may hinder timely AF regeneration. Here, we hypothesized that accelerating PCL degradation kinetics by incorporating a lactide component, known for its rapid degradation, could enhance AF repair. To that aim, PCLA polymers of ε-caprolactone and lactide were synthesised as a copolymer (C-CL90%LA10%) or a blend (B-PCL80%PLA20%). These polymers were electrospun into aligned nanofibrous sheets, which were then assembled into a biomimetic multi-lamellar 3D implant. Cytocompatibility was assessed in vitro using ovine AF cells and ex vivo using a bovine tail disc model. In vitro, PCLA sheets promoted ovine AF cell alignment, proliferation, and expression of type I and II collagen. Ex vivo, the multi-lamellar implant remained in place within a full-thickness (4 mm) annular defect in bovine tail discs for 4 weeks, with initial cell infiltration. Degradation and regenerative potential were then evaluated in an ovine lumbar full-thickness annular defect model at 1 month (n=4) and 6 months (n=10) after implantation. Histological and immunohistochemical analyses revealed substantial cellular infiltration and neo-tissue ingrowth within implant layers, despite implant displacement in some cases. At 1 month, all implants retained their multi-lamellar structure with no visible degradation. By 6 months, the copolymer implants exhibited marked degradation, whereas PCL and blend implants remained structurally intact. Furthermore, ex vivo biomechanical testing revealed comparable flexibility to intact but much lower than non-implanted controls in axial rotation. This study confirms the accelerated degradation kinetics of PCLA copolymers within the disc microenvironment and demonstrates that the multi-lamellar PCLA implant can guide AF tissue repair. Further optimization is needed to enhance implant retention and ensure long-term functional integration.

## Introduction

The intervertebral disc (IVD) is a fibrocartilaginous structure located between vertebrae, acting as a shock absorber and contributing to spine mobility. The IVD achieves its function through three main components: the nucleus pulposus (NP), the annulus fibrosus (AF), and the cartilage endplates (CEPs). The CEPs are thin layers of hyaline cartilage that cover the upper and lower edges of the disc. The CEPs facilitate load distribution and nutrient diffusion across the disc. The NP occupies the core of the disc and is a gelatinous, highly hydrated tissue rich in proteoglycans and type II collagen, conferring compressive resistance. The NP is confined within concentric lamellae of parallel collagen fibers oriented in alternating directions, forming the AF. The peripheral AF layers are rich in type I collagen, while inner layers contain more type II collagen and glycosaminoglycans (GAGs)^1–4^. The AF resists tensile, shear, and compressive strain arising from spinal motion and NP pressurizing ^1,3,4^.

Low back pain (LBP) is the leading cause of disability worldwide, with most people experiencing it at least once in their lifetime. LBP is a major cause of work loss and reduced quality of life. In 2020, it affected 619 million people globally, representing a major socioeconomic burden ^2^. Impairment of AF structure, whether due to trauma or progressive degeneration, can lead to LBP. LBP can arise from discogenic pain, where outer AF lesions become invaded by blood vessels and nociceptive fibers (neo-innervation), or from radicular pain caused by NP herniation through AF tears compressing adjacent nerves^1,5^. When conservative treatment, such as analgesic drugs and physiotherapy, fails, microdiscectomy, the standard surgical treatment for herniation, alleviates symptoms by removing herniated NP material but leaves the AF defect unrepaired^6^. Due to its limited intrinsic healing capacity, AF tears do not regenerate effectively. The repair process typically results in disorganized fibrous repair tissue and incomplete closure of the inner AF region ^1,7^. Consequently, the disc’s structural and mechanical integrity are not restored, placing it at risk of reherniation (25% of the cases^8^) and accelerated degeneration due to impaired mechanical function^6,9,10^.

Therefore, there is a growing need for repair strategies that can restore both the structural integrity and mechanical function of the disc following herniation. Among emerging strategies, implant devices that combine mechanical reinforcement with bioactivity show particular promise for restoring disc structure and function. Currently, the Barricaid™ system, which consists of a woven polyethylene terephthalate (PET) mesh inserted into the AF defect and anchored to the adjacent vertebrae with a titanium screw, is the only device available on the market ^11^. Clinical studies have reported pain relief, preservation of disc height, and reduced reoperation and reherniation rates ^12–14^, with outcomes sustained for up to five years ^15^. However, its invasive implantation method raises concerns about long-term endplate damage, as evidenced by radiography findings, although they did not affect clinical outcomes ^8,15^. Furthermore, the device is only suitable for large AF defects (6-10mm) and does not induce a natural repair process. Therefore, there remains a critical need for treatment strategies that can induce AF regeneration through implants possessing both suitable mechanical properties and a structure that mimics the native AF.

Several studies have evaluated implants made of natural or synthetic polymers with promising results, although none have reached clinical translation. Examples include natural polymers such as collagen^16–26^, chitosan^25,27^, silk^28–33^, as well as synthetic materials such as polycaprolactone (PCL)^34–47^, polyurethane (PU)^23,36,48–51^, and polylactic acid (PLA)^52–59^. These studies have been hindered by the AF’s complex structural organization, limited regenerative capacity, and intricate anatomy. Furthermore, they consistently highlight one of the most significant challenges of AF closure: maintaining proper implant retention under physiological disc loading^60,61^. Therefore, effective annulus fibrosus repair requires materials capable of withstanding the disc’s demanding mechanical environment while guiding functional tissue regeneration.

In our previous work, we used an implant made of PCL, a biocompatible polymer with high mechanical strength that has been widely used in tissue engineering and clinical applications^62–64^. We demonstrated that PCL polymer sheets exhibit uniaxial tensile properties comparable to a single AF lamella ^65^; the Young’s Modulus of a single AF lamella reported in literature is 31-65 MPa ^66^. Furthermore, the PCL-based implant integrated well with the surrounding AF tissue, leading to AF-like neo-tissue formation. However, the slow degradation of PCL (2-4 years ^67,68^) may hinder timely repair tissue deposition and thus, restoration of disc mechanics, increasing the risk of long-term repair failure. Modulation of PCL degradation kinetics can be achieved by incorporating other polymers, such as PLA, to form PCLA copolymers or blends^69,70^. In our previous study, we demonstrated that electrospun aligned PCLA nanofibrous sheets exhibited tensile properties comparable to those of a single AF lamella (Young’s modulus 20–36 MPa) ^71^. However, the effect of polymer degradation kinetics on AF repair in vivo has not yet been explored.

In this study, we investigated the influence of PCLA polymer composition and structure (copolymer vs Blend) on degradation behavior within the injured disc microenvironment. We first determined whether aligned, electrospun PCLA sheets could induce spontaneous organization of ovine AF cells in vitro. Then, we evaluated ex vivo the implantability of a biomimetic multilayer implant in a bovine tail disc. Finally, the regeneration potential of the three implant types (PCL, PCLA copolymer, and PCLA blend) was examined in an ovine lumbar full-thickness annular defect model at 1 and 6 months post-implantation.

## Material and methods

### Polymer sheets synthesis

Polycaprolactone (PCL, M̅n = 80 kgꞏmol⁻¹) was purchased from Sigma-Aldrich. Polymers were synthesized as previously described ^71^. Briefly, Poly(D,L-lactide) (PLA) was synthesized by bulk ring-opening polymerization (ROP) of D,L-lactide using benzyl alcohol and SnOct₂. Poly(D,L-lactide-co-ε-caprolactone) (PCLA, 150 kgꞏmol⁻¹) was synthesized by ROP of ε-caprolactone (CL) and lactide (LA) at 90/10 molar ratios using diethylene glycol as initiator and SnOct₂. After dissolution in dichloromethane (DCM), polymers were purified by precipitation in cold methanol and vacuum drying. Polymer blends were prepared by dissolving commercial PCL and synthesized PLA in Trifluoroethanol (TFE) at an 80/20 CL/LA molar ratio. Polymers are denoted as C-CLxLAy (copolymers) or B-CLxLAy (blends), where x and y represent mol% of caprolactone and lactide. Electrospinning was performed using a Linari Nanotech "Starter Kit-Aligned" from 15–25% (w/v) TFE solutions under 12–14 kV at 1.5 mLꞏh⁻¹. Aligned fibrous sheets (150 ± 30 μm) were collected on an 8 cm rotating wheel (2000 rpm). Then, the samples were sterilized by gamma irradiation at 25 ± 10% kGy (Ionisos, France).

### Multilayer implant preparation

A fibrous layer was first folded four times. Circular discs were then cut from this folded layer using a biopsy punch (4 mm and 2 mm diameter for ex vivo and in vivo implantation, respectively). The discs were stacked to a length of 6mm, then sutured together using a sterile synthetic absorbable suture thread (Peters Surgical OPTIME ET1200(V17)-10/2022). To maintain the structure, a knot was tied on the surface of the external disc. For the ex vivo experiment, an anchoring membrane, or "fixation wing," was added to the implant. This membrane was fabricated by compression molding. Briefly, 1 ± 0.20 g of copolymer C-CL_70%_LA_30%_ was placed between two PTFE sheets, heated to 70 °C, and pressed at 1.5 tons. The resulting films were approximately 400 µm thick and were cut with a scalpel into 10 mm by 5 mm rectangles.

### Cell isolation and culture

Annulus fibrosus (AF) cells were isolated from the lumbar intervertebral discs (IVDs) of three 6-month-old lambs under sterile conditions. Briefly, AF tissue was separated from the rest of the disc and carefully minced. The tissue pieces were rinsed with Hanks’ Balanced Salt Solution (HBSS) before undergoing a multi-step enzymatic digestion at 37 °C. First, the tissue was incubated with 0.05% hyaluronidase (Sigma H4272) for 15 minutes, followed by a 30-minute incubation in 0.2% trypsin (Sigma T9935), and then a final overnight incubation in a 0.2% collagenase solution (Sigma C5138). The resulting cell suspension was then filtered through a 70 µm filter and centrifuged at 1400 rpm for 5 minutes. The cells were seeded at a density of 40.10^3^ cells/cm^2^. Cells were cultured in complete medium composed of Dulbecco’s Modified Eagle Medium (DMEM) high glucose, GlutaMAX™ Supplement, pyruvate (Gibco, 31966-021), supplemented with 10% fetal bovine serum solution (FBS, CVFSCF00-01, Eurobio Scientific), 1 % Penicillin-Streptomycin (10,000 U/mL, Thermo Fischer Scientific: 15140122), 0.1% Amphotericin B (Thermo Fischer Scientific, 15290026). The medium was changed twice a week, and cells were used or passaged when reaching 70%-80% confluency. The cells were reseeded at a density of 5000 cells/cm^2^ and were used up to passage 4 in the subsequent experiments. Fibrous sheets (0.8 x 0.8 mm) of PCL, copolymer, or blend polymers were placed into 24-well plates pre-coated with 2% agarose. The sheets were pre-treated with FBS for 2 hours before being seeded with AF cells at a density of 5000 cells/cm^2^ in complete culture medium supplemented with 0.17 mM ascorbic acid (2-phospho-L-ascorbic acid trisodium salt, Sigma-Aldrich, Gibco, 31966-021). AF cells seeded directly on plastic served as a control. The plates were maintained in a humidified incubator with 5% CO₂ at 37 °C. The medium was changed every 2-3 days.

### EDU labelling

Proliferative cells were labelled using EdU Click-iT™ Plus kit (C10637, Thermo Fisher Scientific). On day 6, cells were incubated with a 10 µM solution of 5-ethynyl-2′-deoxyuridine (EdU) for 24 hours. Following incubation, they were fixed with 4% paraformaldehyde for 15 minutes at room temperature. EdU detection and Hoechst labelling were then performed according to the manufacturer’s specifications. Images were acquired using a fluorescence microscope (Axio Zoom V16). Quantitative analysis was performed using Qupath software (v:0.4.4). (N=1, n=3).

### DNA Quantification

The increase in DNA quantity was evaluated using a Quant-iT™ PicoGreen™ dsDNA Assay Kit (P7589, Thermo Fisher Scientific) according to the manufacturer’s instructions. Fibrous sheets without seeded cells were used as a negative control. The fluorescence signal was measured using a TriStar² S LB 942 multimode reader (Berthold Technologies) with excitation at 480 nm and emission at 520 nm. (N=3, n=3).

### Metabolic activity

The metabolic activity of the seeded cells was quantified using Cell Counting Kit-8 (CCK8, CK04-11, Dojindo) on days 1, 7, 14, following the manufacturer’s instructions. Cells were incubated in a CCK-8 solution for three hours, and then absorbance was measured at 450 nm using a TriStar 2 S LB 942 multimode reader (Berthold Technologies). Fibrous sheets without seeded cells served as a negative control, and the staining solution served as a blank. Optical density values from the samples were normalized to the blank and seeding areas. (N=3, n=3).

### Cell morphology and phenotype

On day 7, implants were fixed with 4% paraformaldehyde for 30 minutes. Then, the cells were permeabilized with 0.5% Triton X-100® and non-specific sites blocked with 4% bovine serum albumin (BSA, Sigma-Aldrich) for 30 minutes. Subsequently, the implants were incubated overnight at 4 °C with an anti-collagen type I primary antibody (1:250 in 1% BSA solution, Ab138492, rabbit monoclonal, Abcam, Cambridge, UK) or an anti-collagen type II antibody (1:250 in 1% BSA, Ab34712, rabbit polyclonal, Abcam). Then, implants were incubated with Alexa Fluor 488 secondary antibody (Goat anti-Rabbit IgG (H+L), 1:500 in 1% BSA solution, Thermo Fisher Scientific) for 1 h at room temperature. The actin cytoskeleton was stained with Phalloidin-iFluor 647 (A22287, Thermo Fisher Scientific) for 1 hour at room temperature. The cell nuclei were stained with Hoechst (4 μg/ml, Thermo Fisher Scientific, 15290026), and images were acquired using a confocal microscope (A1RS, Nikon, France). (N = 1, n =3). Quantitative analyses of directionality were performed using ImageJ® software.

### Scanning electron microscope (SEM)

On days 7 and 14, the implants were fixed with 4% paraformaldehyde for 15 minutes at room temperature, followed by a second fixation with 2.5% (v/v) glutaraldehyde in 0.1 mol/L PBS for 1 hour. The samples were then incubated in 2% (v/v) osmium for 1.5 hours. They were subsequently washed three times with distilled water and dehydrated in an ethanol series with increasing concentrations. The samples were then dried using a gradient of ethanol/Freon washes. Finally, the dried implants were coated with a gold-palladium alloy. Samples were observed with a scanning electron microscope (SEM, Gemini 300, ZEISS) at 2 kV with a 30 µm diaphragm and the SE2 detector positioned in the chamber. (N=1, n=1).

### Ex vivo experiments

#### Disc isolation and culture

Thirteen tail discs were isolated from three female cows (4–6 years old) obtained from Oniris (National Veterinary School of Nantes, France). The animals were euthanized for reasons unrelated to research. The tails were cleaned with a 10% Betadine solution. After the soft tissue was removed, the discs were harvested using a reciprocating saw (Stryker). The harvested discs were initially washed in Betadine, followed by a 15-minute wash in a solution of 3% penicillin-streptomycin. A defect was then created using a 4 mm biopsy punch. Three discs were implanted with PCL implants, three with copolymer implants, and three with blend implants. Additionally, two discs were kept intact as a control, and two discs with empty defects were used as a negative control. Implants were placed in FBS for 1 hour before implantation. The discs were then cultured for 4 weeks in Dulbecco’s Modified Eagle Medium (DMEM) high glucose, supplemented with GlutaMAX™ (Gibco, 31966-021) and pyruvate. The medium was supplemented with 10% FBS, 2% penicillin-streptomycin, 0.1% Amphotericin B, and 0.17 mM ascorbic acid. The medium was changed twice a week.

### In vivo experiments

#### Ethical aspects and animals

A total of 14 female sheep (aged 4-8 years, weight: 50-80 kg, Vendée breed, from GAEC HEAS farm, Ligné F-44850, France) were used for the study. Surgical procedures were performed at the accredited Centre of Research and Preclinical Investigations at the Oniris (National Veterinary School of Nantes). All animal experiments were approved by the French Ministry of Agriculture and the ethics committee of the Région Pays de La Loire (Ethical number APAFIS 47969) and were conducted in accordance with the EU Directive 2010/63/EU. Before the procedure, 20 mL of blood was drawn from the jugular vein, transferred into a sterile polypropylene tube, and allowed to coagulate for 30 minutes at room temperature. The tubes were then centrifuged at 10,000 g for 15 minutes, after which the serum was recovered and stored at -20°C. Before implantation, the implants were incubated in this serum for 1 hour.

### Surgical procedure

All procedures were performed under general anesthesia with jugular venous access. Premedication consisted of a combination of 0.2-0.3 mg/kg of diazepam (not exceeding 20 mg total) (The brand name isn’t needed; the generic name is sufficient), medetomidine 4–6 µg/kg IV (maximum 0.2 mg total), and methadone 0.3–0.5 mg/kg IV (maximum 20 mg total). Anesthesia was induced using ketamine 3–5 mg/kg IV (up to 400 mg) and propofol 0.5–1 mg/kg IV (up to 100 mg). Anesthesia was maintained with 1–3% isoflurane in oxygen at 2 L/min. Intraoperative analgesia was obtained by methadone as per premedication with supplemental 0.03 mg/kg IV boluses as required, together with a continuous rate infusion (CRI) of lidocaine 3 mg/kg/h plus ketamine 0.5 mg/kg/h maintained for the duration of surgery. Postoperative analgesia included meloxicam 0.5 mg/kg IV and buprenorphine 0.01 mg/kg IV administered at the end of surgery, followed by meloxicam 0.5 mg/kg SC once daily on postoperative Days 1-3. Perioperative antimicrobial prophylaxis consisted of long-acting amoxicillin 15 mg/kg SC. All doses were titrated within the specified ranges to achieve the intended clinical effect. The sheep were carefully positioned on their right sides, and their lumbar discs were exposed via a ventrolateral approach. Five IVDs (L1–L2 to L5–L6) were used from each sheep. The IVDs were divided into the following experimental groups (n=14 per group): an intact group, a non-implanted group, and three repaired groups based on the implanted implant material (PCL, copolymer, or blend). A defect was created using a 2 mm biopsy punch on the ventrolateral side of the disc. A nucleotomy was then mimicked by the removal of nucleus pulposus (NP) tissue (80 mg average explant weight) using a surgical rongeur. For the 1-month time point, fixation wings were glued to the adjacent vertebrae using medical-grade cyanoacrylate adhesive (72541-04, Leukoplast), and an external polytetrafluoroethylene (PTFE) patch (GORE® PRECLUDE® Pericardial Membrane, PCM102, GORE) was also glued with cyanoacrylate adhesive to the adjacent vertebral bodies. For the 6-month time point, the implant was secured in place by suturing the AF defect using ETHILON thread (polyamide 5-0, PA600444, ETHICON). At the 1-month and 6-month time points, the sheep were sacrificed under general anesthesia via the jugular vein with sodium pentobarbital 400 mg/mL, 20 mL IV. Four sheep were sacrificed at one month, and their discs were processed for histological analysis. Ten sheep were sacrificed after six months; discs from four of these sheep were processed for histological analysis, while the lumbar spines from the remaining six sheep were processed for biomechanical analysis.

### X-ray imaging and MRI

MRI and X-ray imaging were performed preoperatively and at 1, 3, and 6 months. X-ray imaging was performed, as previously described^72^, using a radiography machine (Convix 80® generator and Universix 120 table) from Picker International (Uniontown, Ohio, USA). Coronal and sagittal plane radiographs of the sheep lumbar spines were acquired with a collimator-to-film distance of 100 cm, an exposure of 30-60 mAs, and a penetration power of 70-80 kVp. MRI of the entire lumbar sheep spines was performed as previously described^72^ using a 1.5 T MRI scanner (Magnetom Essenza®, Siemens Medical Solutions, Erlangen, Germany) with a standard spine coil to obtain T2-weighted images (TE: 86 ms, TR: 3000 ms; slice thickness: 3 mm) and T1-weighted images (TE: 12 ms, TR: 322 ms; slice thickness: 3 mm). These sequences were followed by the acquisition of sagittal (1) variable flip-angle T1 mapping sequence (repetition time: 15 ms; echo time: 1.7 ms; flip angle: 5°/26°; thickness: 3 mm; field of view: 256 × 264; bandwidth: 280; pixel size: 1 × 1 mm), (2) multi-echo T2 mapping (repetition time: 1510 ms; echo time: 13.8/27.6/41.4/55.2/69 ms; flip angle: 180°; thickness: 3 mm; field of view: 256 × 264; bandwidth: 225; pixel size: 1 × 1 mm), and (3) multi-echo T2* mapping (repetition time: 428 ms; echo time: 4.35/11.83/19.31/26.79/34.27 ms; flip angle: 69°; thickness: 3 mm; field of view: 256 × 264; bandwidth: 260; pixel size: 1 × 1 mm). The images were analyzed using OsiriX software® (3.9, Osirix Foundation, Geneva, Switzerland). Each intervertebral disc (IVD) received a degeneration grade on mid-sagittal T2-weighted images using the 5-point system described by Pfirrmann et al. (PMID: 11568697). In addition, T2, T1 and T2* relaxation times for each NP were subsequently quantified on the mapping sequences by placing a central circular ROI on each NP.

### Histological analysis

The samples were fixed with 4% paraformaldehyde at 4 °C for one week. They were then decalcified for three weeks in Shandon TBD-2™ Decalcifier (12607926, Thermo Fisher Scientific). Following decalcification, the discs were fixed again in PFA for three days. The discs were flash frozen in an isopentane/liquid nitrogen bath and embedded in Super Cryoembedding Medium (SCEM) (Section Lab, Hiroshima, Japan). The embedded samples were frozen again in the isopentane/liquid nitrogen bath until the SCEM solidified. The frozen samples were sectioned at -30 °C in the axial orientation into 10 µm sections using a cryostat (CryoStar NX70®, Thermo Fisher Scientific, Waltham, Massachusetts, USA) and collected using adhesive films (Cryofilm type IIc, SECTION-LAB Co. Ltd., Japan). Following standard protocols, the sections were stained with either Hematoxylin/Eosin/Safran (HES) or Alcian blue.

For immunostaining, sections were incubated with 0.1% trypsin (T9935, Sigma-Aldrich) for 30 minutes at 37 °C, followed by a freshly prepared 3% H₂O₂ solution to inactivate endogenous peroxidases. The sections were then blocked with 4% BSA for 30 minutes before being incubated overnight at 4 °C with the primary antibodies: anti-collagen type I (1:250 in 4% BSA, Ab138492, rabbit monoclonal, Abcam) and anti-collagen type II (1:400 in 4% BSA, MP8631713, mouse monoclonal, MP Biomedicals). Biotinylated goat anti-rabbit and goat anti-mouse secondary antibodies (E432 and E433, respectively, 1:300 dilution, Dako, Agilent Technologies) were used for 1 hour at room temperature. Following secondary antibody incubation, the sections were incubated with horseradish peroxidase-conjugated streptavidin (1:400, P0397, Agilent Technologies) for 45 minutes. Antibody binding was visualized using diaminobenzidine (DAB, K3468, Agilent Technologies) as the HRP substrate, and the sections were counterstained with hematoxylin for 1 minute. All stained sections were observed using a slide scanner (Nanozoomer S360, Hamamatsu Photonics), and images were analyzed with NDP view2 software® (Hamamatsu Photonics).

### Ex vivo biomechanical testing

For flexibility testing, n = 6 lumbar spine specimens (L1-L6) were harvested and stored at -20 °C. In each specimen, the L1-L2 disc had been non-implanted control (n = 6) and the L5-L6 disc had been kept intact (n = 6), whereas the L2-L3, L3-L4, and L4-L5 discs had received randomized treatment with PCL (n = 6), copolymer (n = 6), and blend (n = 6) fibrous scaffolds, respectively. Following thawing at 5 °C for 12 h, the specimens were prepared by carefully removing any muscle and fat tissue and by preserving all passively stabilizing anatomical structures (vertebrae, intervertebral discs, and ligaments). For biomechanical testing, the L1 and L6 vertebrae were half-embedded in PMMA (Technovit 3040, Heraeus Kulzer, Wehrheim, Germany) and equipped with metal flanges for fixation in the testing device. The specimens were loaded with displacement-controlled (1 °/s) pure moments of 7.5 Nm ^73^ in flexion/extension, lateral bending, and axial rotation using a well-established spine tester ^74^. During load application, optical motion tracking of each vertebra was performed by means of three retroreflective markers mounted on each vertebra and twelve infrared cameras (Vicon MX13+, Vicon Motion Systems Ltd., Oxford, UK). Loading was performed for 3.5 cycles, of which the third full load cycle was used for evaluation of range of motion (ROM) and neutral zone (NZ) to minimize viscoelastic effects ^75^.

### Statistical analysis

All of the data are presented as means ± standard deviation. The statistical analyses were performed using GraphPad Prism 5.0® software (GraphPad, San Diego, California, USA). Statistical significance was determined using two-way ANOVA followed by Tukey’s post-test for multiple group comparisons with different variables (CCK8, Picogreen, MRI), one-way ANOVA followed by Tukey’s post-test for multiple group comparisons with one variable (EdU, Pfirmann).) or pairwise Kruskal-Wallis test with Dunn-Bonferroni post-hoc correction for multiple groups with low sample sizes (ROM, NZ). Statistical significance was set at p < 0.05. Unless otherwise stated, the experiments were repeated at least three times.

## Results

### Fibrous scaffolds (PCL, Copolymer, or Blend) guide the alignment and proliferation of ovine AF cells while maintaining collagen type I and type II expression

First, we aimed at assessing the cytocompatibility of the polymers with AF cells in vitro. For this purpose, ovine AF cells were seeded (5,000 cells/cm²) onto the fibrous sheets of the polymers (PCL, Copolymer, or Blend) and cultured for up to 14 days. Cells cultured on plastic (2D) were used as a control. AF cells colonized the surface of the sheets as identified by fluorescence microscopy imaging of F-actin staining after 7 days of culture (Figure 1A). Cells were uniformly distributed on the surface and exhibited a fusiform shape, compared to the irregular morphology of 2D cultured cells. The alignment of AF cells was investigated by analyzing actin filament orientation using the Directionality plugin in ImageJ (Figure 1B and Table S1). For convenience, 0° was considered parallel to the horizontal axis. Cells cultured on plastic did not exhibit any specific orientation, resulting in a uniform histogram. Whereas cells cultured on the sheets displayed a bell-shaped histogram with an alignment peak at 0°, indicating that actin filaments, and thus the cells, were predominantly aligned parallel to each other.

**Figure 1.**
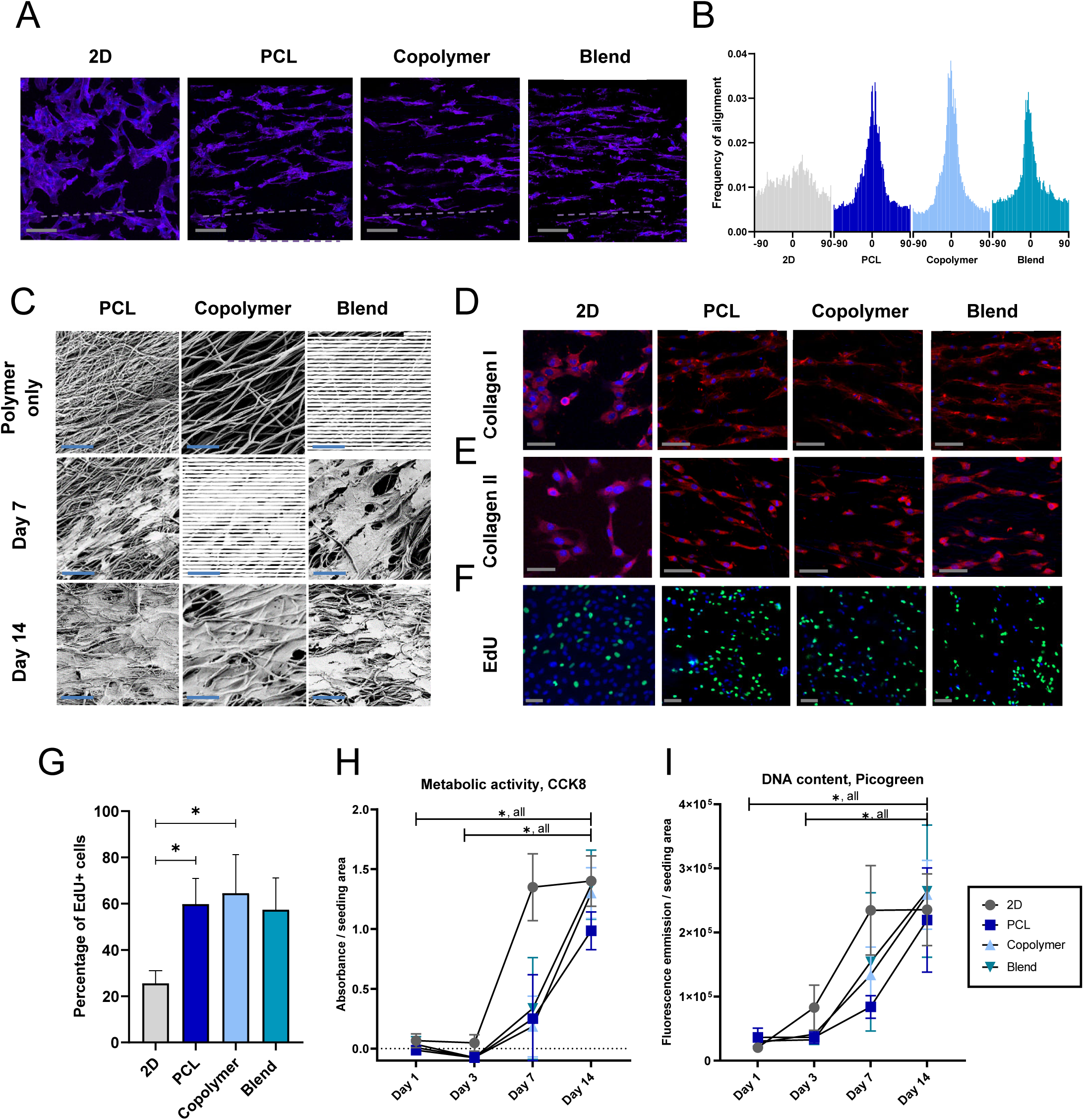
The polymers sheets promoted AF cell’s alignment and proliferation while maintaining the expression of collagen type I and type II. AF cells (5, 000 cells/cm^2^) were seeded onto the fibrous sheets and cultured up to 14 days. Cells cultured in 2D were used as control. (**A**) Representative fluorescence microscopy images of F-actin staining (Phalloidin) after 7 days of culture. Scale bar: 100 µm. F-actin: magenta, nuclei: blue. Dotted line indicates the fiber orientation. (**B**) Frequency of AF cells alignment after 7 days of culture (based on analysis of F-actin filament alignment). N = 1, n = 3, ; 0° considered parallel to the horizontal axis. (**C**) SEM images of the fibrous sheets seeded with AF cells. Scale bar: 20 µm (**D** and **E**) Representative immunostainings images of AF cells stained for collagen Type I (**D**) and II (**E**) at day 7. Scale bar: 50 µm. Collagen: red, nuclei: Blue. (**F**) EdU labeling for proliferative cells at day 7. EdU: green, nuclei: blue. Scale bar: 50 µm. (**G**) Quantification of EdU^+^ cells, presented relative to the total number of cells (N = 1, n = 3). (**H**) AF cells metabolic activity analysis by CCK-8 assay (N = 3, n = 3). (**I**) DNA quantification by PicoGreen™ dsDNA assay (N = 3, n = 3). All data are represented as mean ± standard deviation. Comparisons among groups were performed using one-way (G) or two-way (H&I) ANOVA, followed by a Tukey’s test. ∗ = statistically significant, p ≤ 0.05. PCL: polycaprolactone, Copolymer: C-CL_90%_LA_10%_, Blend: B-PCL_80%_PLA_20%_, SEM: scanning electron microscopy.

AF cells cultured on the sheets had an alignment score close to 1, whereas those cultured in 2D had a score of only 0.46 (Table S1). The effect of the substrate was further confirmed by SEM images (Figure 2C). Furthermore, synthesis of type I and II collagens, key markers of AF cells, was analyzed by immunostaining (Fig. 1D-E). In all conditions, AF cells were similarly positive for both markers, indicating that the polymer type or surface did not impede collagen type I or type II expression. AF cell proliferation on the polymer sheets was analyzed using 3 complementary assays: Click EdU, CCK8, and PicoGreen (Figure 1G-I). EdU-positive cells were detected by fluorescence staining after 24 hours of incubation with the modified thymidine analog EdU (Figure 1F). The percentage of EdU-positive cells relative to the total number of Hoechst-positive cells (Figure 1G) was similar for the three polymer sheets and significantly higher than for the 2D-cultured cells for PCL and copolymer. Cellular metabolic activity measured using the CCK8 assay (Figure 1H) showed an increase in the metabolic activity of AF cells between day 1 and day 14. Interestingly, cells cultured on the polymer sheets exhibited a delayed increase in metabolic activity between days 1 and 7, followed by a sharp increase between days 7 and 14. On day 7, 2D-cultured cells had significantly higher metabolic activity than polymer sheets, whereas on day 14, all groups displayed a similar activity. Similarly, the PicoGreen assay showed an increase in absorbance between day 1 and 14, with a delayed increase between day 1 and day 7 for cells on polymer sheets, indicating an increase in DNA quantity over time and thus cell proliferation. Together, these data suggest that the polymer sheets not only supported AF cell attachment and viability but also promoted proliferation and alignment without altering the expression of tissue markers.

**Figure 2.**
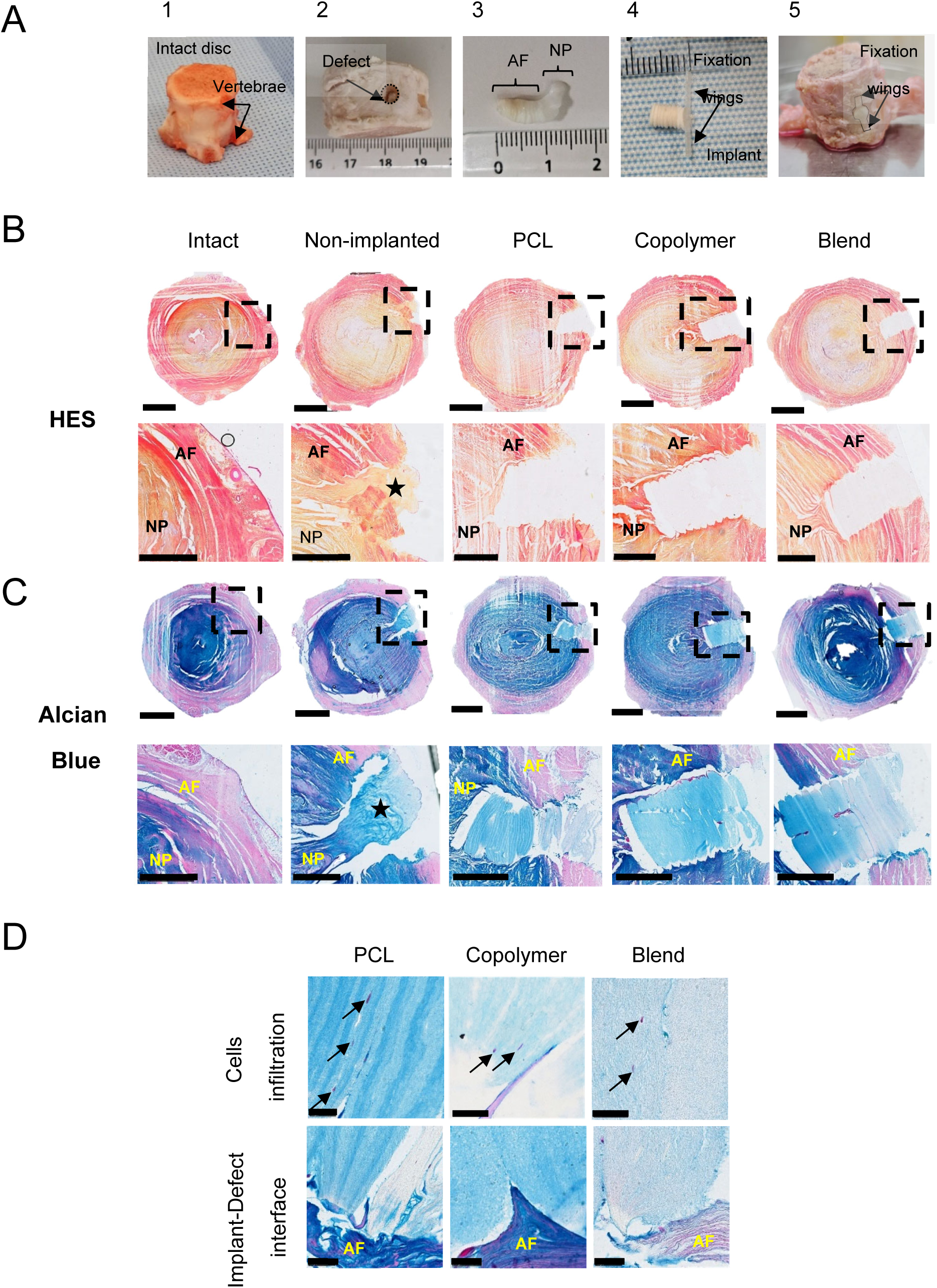
Ex vivo implantation of a multi-lamellar AF-mimicking scaffold in a bovine tail disc model. A full-thickness annular defect with partial nucleotomy was created using a 4 mm biopsy punch, and multilayer implants (PCL, Copolymer or Blend) were placed and secured in the defect by gluing the fixation wings to the adjacent vertebrae (n = 3). The discs were maintained in culture for 1 month. Discs were, cryo-sectioned (10 µm-thick sections), and histochemically stained with Hematoxylin/Eosin/Safran (HES) or Alcian Blue. (**A**) Macroscopic images of: (**A1**) the intact disc, (**A2**) disc with empty defect, (**A3**) recovered AF/NP, (**A4**) implant, (A5) a disc with implant inside the defect. (**B** and **C**) HES (**B**) and Alcian Blue (**C**) staining of the discs after 1 month of culture. Dotted lines indicate the location of the high magnification image. Star: indicates NP extrusion. Scale bar: 5 mm for full-disc images (top row), 2.5 mm for high magnification images (bottom row). (**D**) Alcian Blue stained sections showing cell infiltration into the implant (top, black arrows) and implant-defect interface (bottom). Scale bar: 100 μm. PCL: polycaprolactone, Copolymer: C-CL_90%_LA_10%_, Blend: B-PCL_80%_PLA_20%_, HES: Hematoxylin/eosin/safran.

### Ex vivo implantation of a multi-lamellar AF-mimicking implant in a bovine tail disc model

Next, we evaluated the feasibility of implanting the polymer sheets as a repair device and tested them ex vivo. For that, an implant mimicking the AF multi-lamellar structure was constructed by stacking several layers of polymer sheets (2 mm in diameter) until reaching a length of 6 mm, approximately equal to the average AF thickness of an ovine disc. An anchoring system of “fixation wings” was incorporated, consisting of a flexible rectangular film (copolymer C-CL_70%_LA_30%_), designed to be glued to the adjacent vertebrae. The implantability of the scaffold was tested using a bovine tail disc model (Figure 2A). After the soft tissue was removed, the discs were harvested using a reciprocating saw (Figure 2A1). Then, discs were either kept intact or a full-thickness AF defect was created using a 4 mm biopsy punch (Figure 2A2), followed by partial nucleotomy (Figure 2A3). The implant (Figure 2A4) was inserted into the defect, and the fixation wings were glued to the adjacent vertebrae with medical-grade cyanoacrylate glue (Figure 2A5). Discs with the defect left empty served as the non-implanted controls. Discs were cultured for 1 month and checked visually regularly. No signs of implant displacement were noted. After fixation, discs were carefully processed for cryo-embedding and sectioning. Disc sections were histologically stained with HES and Alcian Blue. HES stains collagen fibers in orange, whereas Alcian blue stains GAGs in blue. These stainings showed no apparent alteration of the disc’s structure in the intact group (Figure 2B, C), including normal cellularity, organized AF lamellae, and intense Alcian blue staining in the NP region, indicating the presence of proteoglycans. In the non-implanted discs, the defect remained open with NP expansion toward the defect. Outside the defect area, no apparent changes in the disc structure were observed. In contrast, in the experimental groups, the implants could be readily identified in all the sections. The implants filled the defect spanning the AF depth, remained in place for 1 month, and no NP expansion was observed.

Furthermore, the implants’ multi-lamellar structure remained visually intact across all polymers, with no signs of delamination. Finally, it is worth noting that the implant was in close contact with the adjacent AF tissue (Figure 2D). Initial cell infiltration at the tissue-implant interface was observed, with cells identified between the inner layers of the implant (Figure 2D, black arrows). These results demonstrate that the multi-lamellar implant can be successfully implanted ex vivo, prevent NP expansion, and support cell infiltration.

### The implant’s multi-lamellar structure was maintained in vivo after 1 month, with ingrowth of AF-like tissue between implant layers

After demonstrating feasibility ex vivo, we next tested whether the implants could promote AF repair in vivo. Specifically, we evaluated whether the degradation behaviors observed in vitro^71^ were reproduced in vivo and whether they influenced AF repair, using an ovine IVD injury model at 1 and 6 months. The model involved creating a full-thickness annular defect, followed by partial NP removal. Lumbar discs were exposed via an anterolateral approach (Figure S1A–C), and a full-thickness AF defect was created using a 2 mm biopsy punch (Figure S1D). The implant (Figure S1E) was inserted into the defect (Figure S1F) and secured in place using two methods, depending on the time point. For the 1-month study, fixation wings and a PTFE patch glued to the adjacent vertebrae were used (Figure S1G–H). This method was found to be too cumbersome from a surgical standpoint, and for the 6-month study, the AF defect was stitched using a continuous suture with 2 passages to maintain the implant in place (Figure S1I).

In the 1^st^ study, X-ray and MRI were obtained preoperatively and at 1 month to assess disc height and NP hydration. No signs of disc degeneration were detected preoperatively, with all discs having a Pfirrmann score of 1. No surgery-related complications were observed at any time point (Figure S2A and B). At 1 month, one disc with a blend implant had a Pfirrmann score of 3, although there was no significant difference between the groups (Figure S2C). Quantitative analysis confirmed that disc height and NP hydration (T2 weighted signal intensity) were maintained across all groups, including implanted, intact, and non-implanted discs (Figure S2D and E). Histological staining of disc sections with HES and Alcian Blue showed no apparent alteration of the disc’s structure in the intact group, including normal cellularity, organized AF lamellae, and intense Alcian blue staining in the NP region (Figure 3A and B). In the non-implanted control, no NP expansion was observed. However, the defect was only partially closed, with unorganized fibrotic tissue, and numerous vascular ingrowths were identified, each containing a lumen filled with red blood cells (bright pink/red dots). In the implanted groups, partial or complete implant displacement occurred (Table S2). Two implants (one Blend and one Copolymer) were not found in the defect area. In addition, three implants (one from each polymer type) were found in two connected pieces. Nevertheless, the implants maintained their multi-lamellar structure without signs of delamination or degradation (Figure 3A and B). All implants integrated well with the surrounding AF tissue, with continuous tissue at the implant margin. A foreign body reaction was observed outside of the discs, most probably due to the presence of the glue material (Figure S3). Homogeneous cell infiltration and de novo tissue formation were detected inside the implants, both between and within the polymer layers, with minimal vascular ingrowth (Figure 3A and B). Outside the defect area, no apparent changes in the disc structure were observed. Immunostaining revealed that the de novo tissue in the non-implanted group was positive only for type I collagen. In contrast, tissue ingrowth within the implant layers was positive for both type I and type II collagens (Figure 4). Despite partial implant displacement, these findings suggest that the implant supported AF like tissue ingrowth and integrated with the surrounding AF tissue at 1 month.

**Figure 3.**
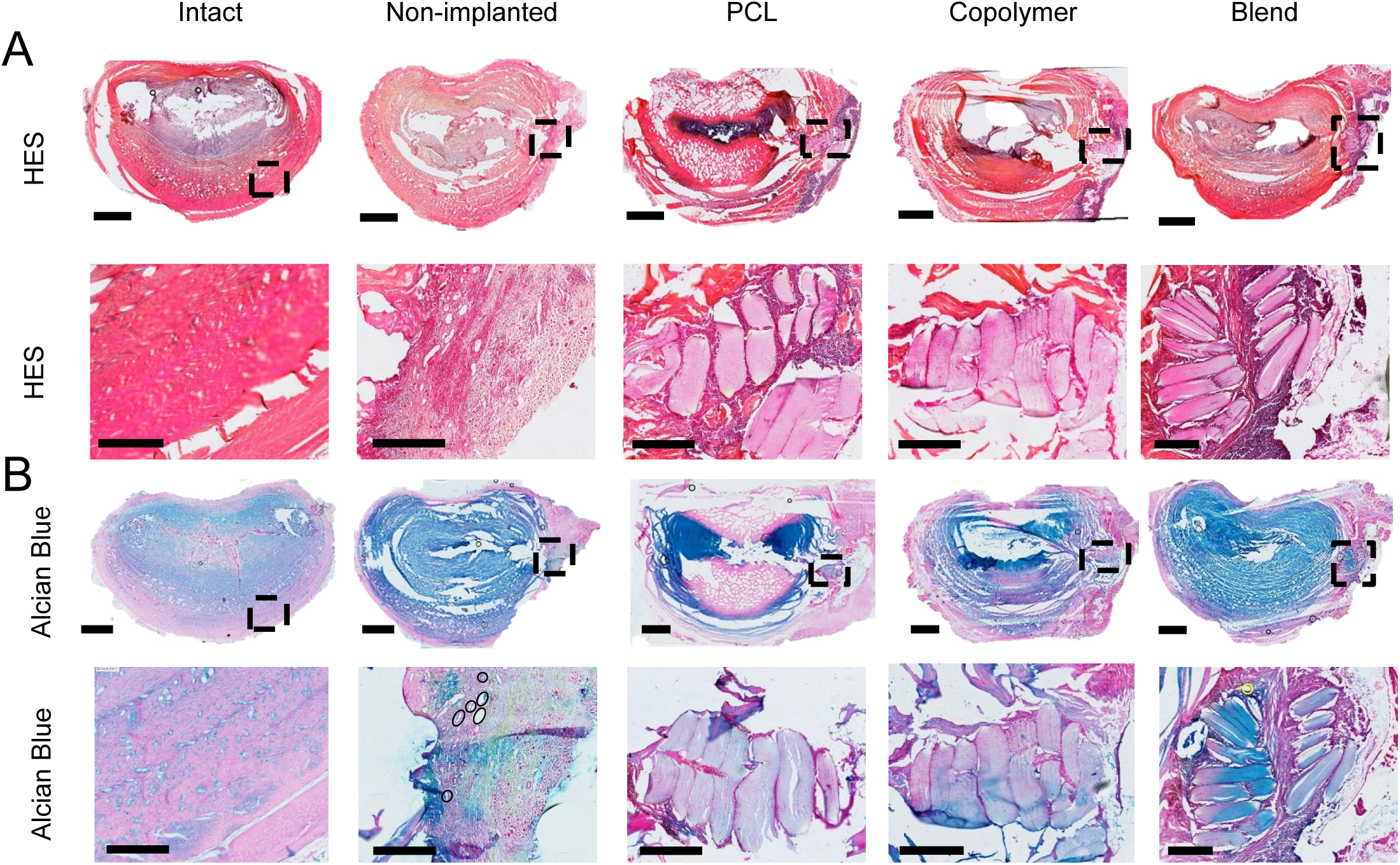
The implant’s multi-lamellar structure was maintained after one month in vivo. A full-thickness annular defect with partial nucleotomy was created using 2 mm biopsy punch, and multilayer implants were placed and secured in the defect by gluing the fixation wings to the adjacent vertebrae. Five IVDs (L1–L2 to L5–L6) were used from each sheep. The IVDs were divided into the following experimental groups (n=4 per group): an intact group, a non-implanted group, and three repaired groups based on the implant material (PCL, copolymer, or blend). Discs were, cryo-sectioned (10 µm-thick sections), and histochemically stained with Hematoxylin/Eosin/Safran (HES) and Alcian Blue. (**A** and **B**) representative images of HES (**A**) and Alcian Blue (**B**) stained section of the disc. Circles identify blood vessels. Dotted lines indicate the location of the high magnification image. Scale bar: 5 mm for full disc images top row, 2.5 mm for high magnification images bottom row). PCL: polycaprolactone, Copolymer: C-CL_90%_LA_10%_, Blend: B-PCL_80%_PLA_20%_, HES: Hematoxylin/Eosin/Safran.

**Figure 4.**
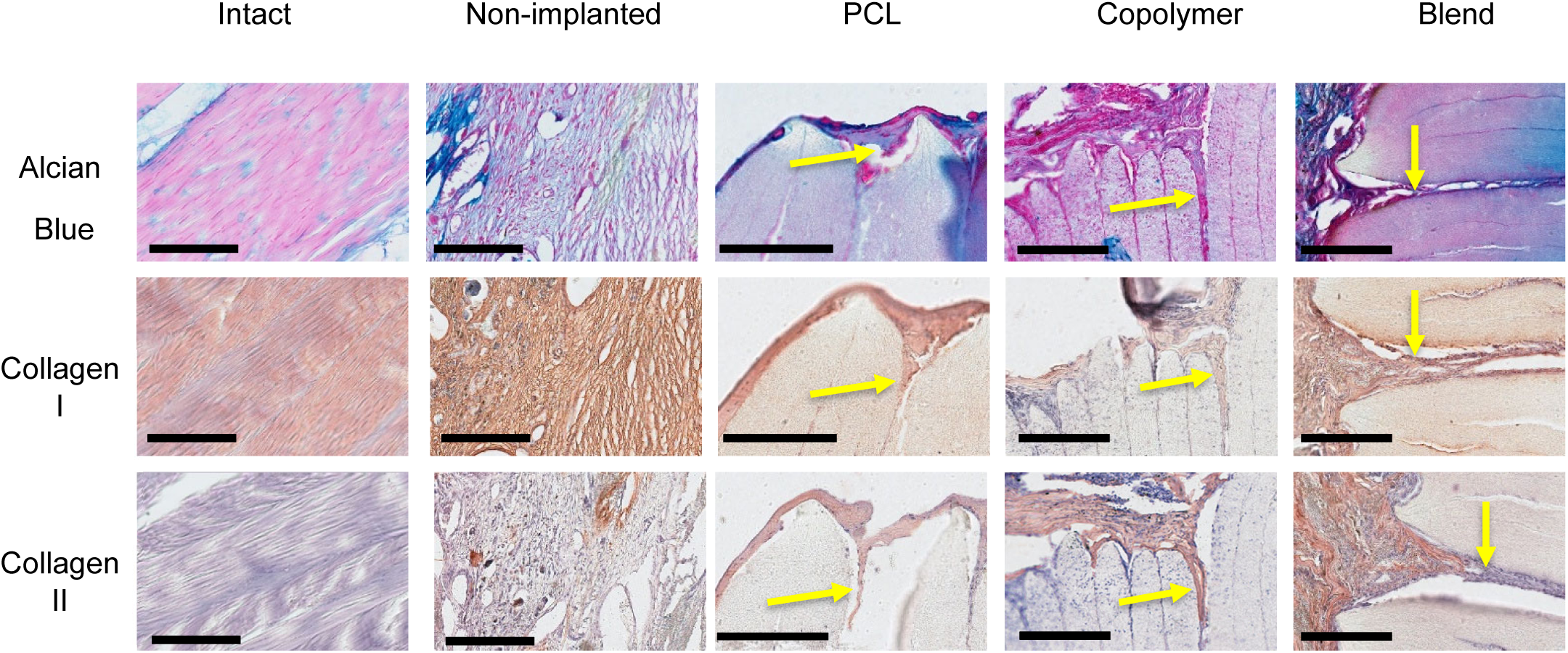
Ingrowth of AF-like tissue between the implant layers. Five groups (n = 4 per group) were compared: an intact group, a non-implanted group, and three repaired groups based on the implant material (PCL, copolymer, or blend). Disc sections were either histochemically stained with Alcian Blue (top row) or immunostained for type I (middle row) and type II (bottom row) collagen. Arrows indicate the tissue ingrowth between implant layers. Scale bar: 250 μm. PCL: polycaprolactone, Copolymer: C-CL_90%_LA_10%_, Blend: B-PCL_80%_PLA_20%_,

### Copolymer implants degraded after 6 months in vivo, while PCL and Blend maintained their structure

We next evaluated the long-term repair efficacy and implant degradation at 6 months. Due to implant displacement and the foreign body reaction observed at 1 month, we modified the implantation strategy for the 6-month study by suturing the AF defect over the implant rather than using glue (Figure S1I).

X-ray and MRI were acquired preoperatively and at 1, 3, and 6 months to monitor disc height and NP hydration. No signs of disc degeneration were detected before surgery, and no surgery-related complications occurred (Figure S4A and B). All discs had a Pfirrmann score of 1-2 preoperatively. Two discs (one with a copolymer implant and one non-implanted control) had a Pfirrmann score that increased from 1 preoperatively to 3 at 6 months (Figure S4C), with no significant differences between groups at each time point. Quantitative analysis confirmed that disc height decreased significantly over time in the non-implanted and Blend groups (Figure S4D), but was preserved in the Intact, PCL, and copolymer groups. Similarly, NP hydration measured by T2 signal intensity showed a delayed loss of hydration in the implanted groups compared with non-implanted controls (Figure S4E). Ex vivo biomechanical testing revealed that none of the discs treated with the three implants (PCL, copolymer, blend) exhibited significant differences in ROM and NZ compared to the intact discs. However, relative to the non-implanted group, discs treated with PCL implants showed significantly lower left and right axial rotation flexibility. Overall, PCL, copolymer, and blend implant-treated discs exhibited similar ROM and NZ compared to the intact discs in flexion/extension and lateral bending, whereas non-implanted discs showed more than twice the ROM compared to the other groups (Figure S5). Histological staining of disc sections with HES and Alcian Blue showed a preservation of overall disc structure in the intact group, including normal cellularity, organized AF lamellae, and intense Alcian blue staining in the NP region (Figure 5A and B). In the non-implanted control, the defect was partially closed with unorganized fibrotic tissue, whereas the inner AF region remained unrepaired. Vascular ingrowths were found in the defect area. In addition, positive Alcian blue staining and round cells within lacunae were observed in the defect area, indicating NP expansion toward the defect. In the implanted groups, no pronounced foreign body reaction was observed within or around the implant where no glue was used, contrary to what was observed in the 1-month study. NP expansion toward the defect area was also observed in the three implanted groups. Furthermore, partial or complete implant displacement occurred across all conditions (Table S3).

**Figure 5.**
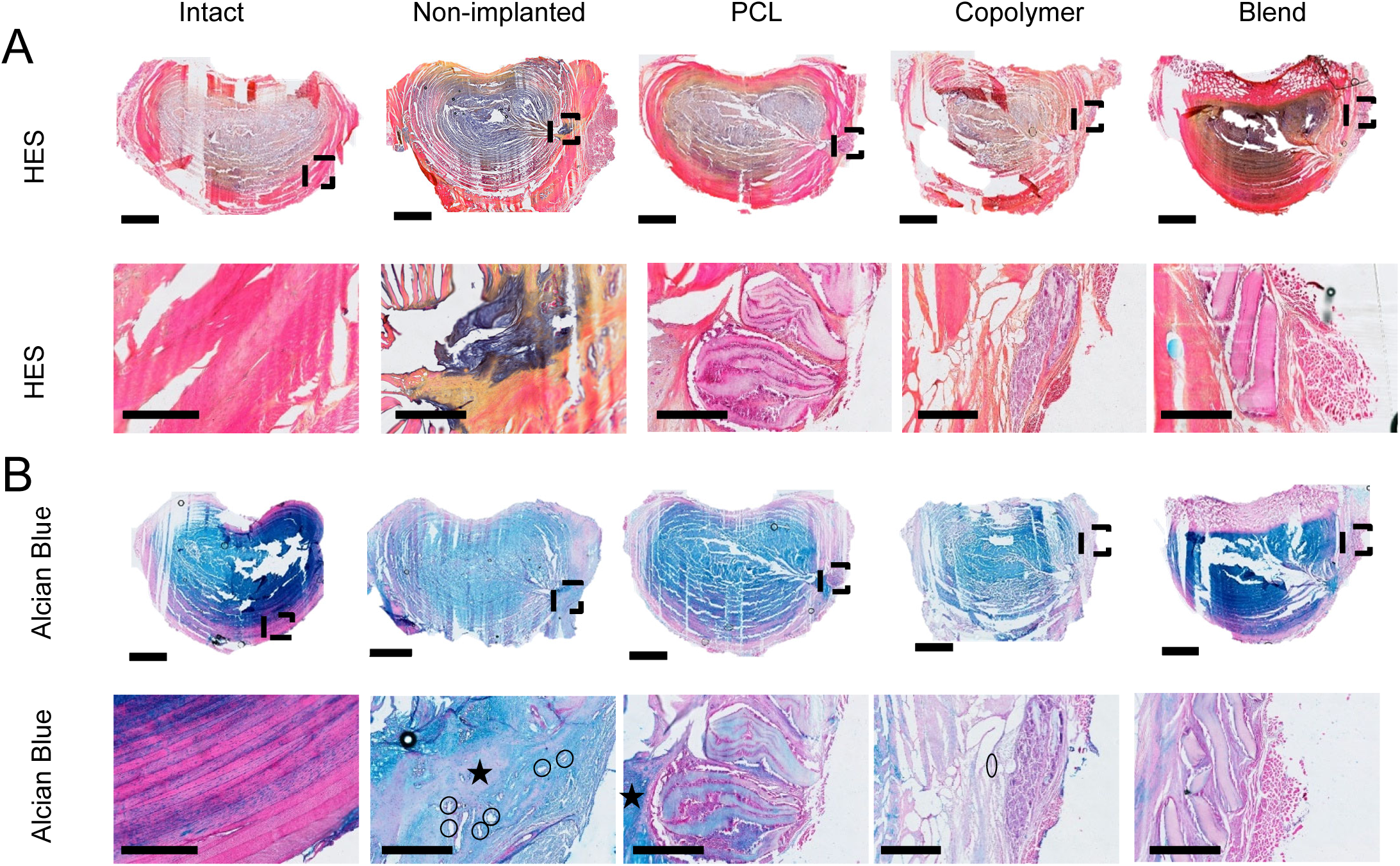
Copolymer implants degraded after 6 months in vivo, while PCL and Blend maintained their structure. A full-thickness annular defect with partial nucleotomy were created using 2 mm biopsy punch, and a multilayer implant was placed and secured in the defect by suturing the AF defect. Five discs (L1–L2 to L5–L6) were used from each sheep. The discs were divided into the following experimental groups (n = 4 per group): an intact group, a non-implanted group, and three repaired groups based on the implant material (PCL, copolymer, or blend). Discs were cryo-sectioned (10 µm-thick sections), and histochemically stained with Hematoxylin/Eosin/Safran (HES) and Alcian Blue. (**A** and **B**) Representative images of HES (**A**) and Alcian Blue (**B**) stained sections of the discs. Circles identify blood vessels. Star: indicates NP extrusion. Dotted lines indicate the location of the high magnification image. Scale bar: 5 mm for full disc images top row, 1 mm for high magnification images bottom row). PCL: polycaprolactone, Copolymer: C-CL_90%_LA_10%_, Blend: B-PCL_80%_PLA_20%_, HES: hematoxylin/eosin/safran.

Interestingly, the PCL and Blend implants could be easily identified on histological sections of 9 out of 10 discs with the lamellae present and easily distinguishable (Figures 5A and B). By contrast, only 4 of 10 copolymer implants were found at 6 months (Table S3), and these were largely degraded, with only small remnants of the layered structure remaining (Figures 5A and B). All implants integrated well with the surrounding AF tissue, with continuous tissue at the implant margin. Compared to the 1-month time point, greater cell infiltration and tissue ingrowth were observed within the PCL and Blend implants, with minimal vascular ingrowth. (Figure 5A and B). For Copolymer implants, AF-like tissue surrounded the degraded implant, while dense cellular infiltration was observed within the residual layers. Immunostaining revealed that the de novo tissue in both the non-implanted group and the implanted groups was positive for type I collagen and negative for type II collagen (Figure 6). Overall, Copolymer implants degraded faster than PCL or Blend in vivo, and this accelerated degradation did not impact AF repair outcomes.

**Figure 6.**
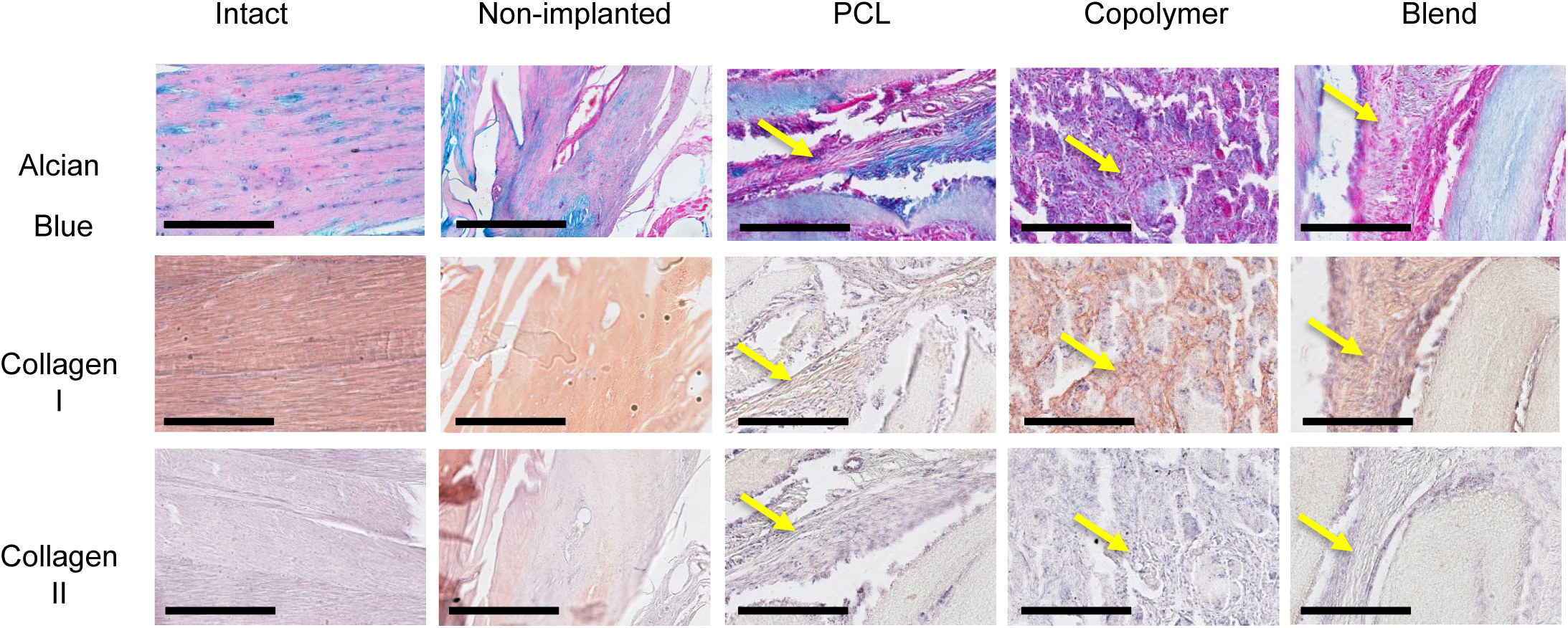
Neo tissue formation between the implant layers. Five groups (n = 4 per group) were compared: an intact group, a non-implanted group, and three repaired groups based on the implant material (PCL, copolymer, or blend). Disc sections were either histochemically stained with Alcian Blue (top row) or immunostained for type I (middle row) and type II (bottom row) collagen. Arrows indicate the tissue ingrowth between implant layers. Scale bar : 250 μm. PCL: polycaprolactone, Copolymer: C-CL_90%_LA_10%_, Blend: B-PCL_80%_PLA_20%_,

## Discussion

In this study, we evaluated a bio-inspired multi-lamellar implant composed of oriented nanofibers with tunable mechanical properties, an anchoring system, and enhanced biodegradability for annulus fibrosus regeneration. In vitro, the implant guided AF cell alignment, proliferation, and maintenance of key phenotypic markers (type I and type II collagens). Ex vivo and in vivo experiments demonstrated the feasibility of implantation and implant integration with surrounding AF tissue, as well as cell infiltration and neo-tissue formation within the implant layers. We further revealed that the polymer structure influenced the in vivo degradation kinetics, with the copolymer implants degrading significantly faster than the PCL or blend counterparts.

The choice of LA percentage is based on our previous in vitro degradation study, which showed that the copolymer (C-CL90%LA10%) and the Blend (B-PCL_80%_PLA_20%_) exhibited comparable molecular weight loss in vitro over six months^71^. While the degradation kinetics of PCLA polymers have been reported to be enhanced by LA incorporation ^76,77^, few studies have evaluated their in vivo degradation behavior. To our knowledge, our study is the first to assess the long-term degradation kinetics of PCLA polymers within the intervertebral disc microenvironment.

We first showed that the polymer type did not affect AF cell morphology, collagen type I and II expression, or proliferation in vitro. As specific phenotypic markers of AF cells are still lacking, we assessed the expression of key ECM proteins (type I and type II collagen). AF cells cultured on the electrospun polymer sheets expressed both type I and type II collagen. AF cells constitute a heterogeneous population of cells with zonal variation within the disc. While outer AF cells are fibroblastic and express a high level of type I collagen, inner AF cells are more chondrocyte-like, expressing type II collagen. This zonal diversity likely explains the dual collagen expression observed in our cells. No evident ECM deposition was detected in vitro, possibly due to the relatively short culture duration (14 days) and/or the diffusion of secreted ECM molecules into the culture medium, consistent with previous reports using electrospun polymer sheets^78,79^. However, matrix accumulation has been observed using compact multilayered electrospun scaffolds^80^. Furthermore, there is no consensus on optimal culture medium formulation for AF cells^81^. The medium used in our study might have lacked essential growth factors such as TGF-β3, which has been shown to promote ECM deposition by AF cells^78^. Alternatively, employing the molecular crowding technique^82^, by adding macromolecules to increase solution density and mimic the in vivo microenvironment, could enhance ECM deposition in vitro. Proliferation analysis (CCK-8, PicoGreen, and EdU assays) revealed an initial slower cell growth on the polymer sheets compared to 2D culture, likely due to lower seeding efficiency. Nevertheless, by day 14, absorbance levels (CCK8 and picogreen assays) were comparable. On the other hand, a lower proportion of EdU-positive cells was observed in 2D cultures, likely reflecting contact inhibition at confluency, whereas cells on the fibrous sheets continued to proliferate on the available surface.

The fibrous sheets were then stacked into a multilayer implant to replicate the multi-lamellar structure of AF tissue and provide more space for cellular infiltration and layered matrix deposition ^78^. Electrospinning technology enabled the fabrication of aligned fibrous sheets mimicking collagen orientation within single AF lamellae^5,83,84^. Aligned fibers were shown to provide contact guidance cues for AF cells and to enhance the mechanical properties of the repair tissue compared to random fibers ^78,80,85–87^. The feasibility of implantation was first assessed using an ex vivo bovine disc model. Implants remained within the defect and maintained contact with surrounding tissue. Cell infiltration was minimal, consistent with previous studies using similar setups ^88^. This limited infiltration likely reflects the absence of mechanical stimulation and the low intrinsic repair capacity of the disc. Future work should evaluate the effect of chemotactic factors ^89^ or dynamic loading ^90^ on enhancing cellular migration into the implant. Furthermore, in non-implanted ex vivo discs under unconfined conditions without mechanical loading, NP bulging toward the defect area was likely due to tissue swelling, given the GAG-rich, hydrophilic nature of NP ^5,91^. We then conducted an in vivo study to evaluate the repair potential of our implant. A large-animal ovine model was selected due to its similarity to human IVD anatomy, composition, and biomechanical properties ^73,92^. In addition, ovine discs exhibit age-related degeneration, characterized by the progressive disappearance of notochordal cells, similarly to what is observed in human discs ^93^. Together, these characteristics make them clinically relevant for translational AF repair studies ^73,92–94^. In this study, a ventrolateral surgical approach was selected to provide safe access to multiple lumbar discs per animal, bypassing the nerves and posterior bone structures. Compared with the posterior standard discectomy surgical approach, the ventrolateral approach reduces postoperative pain, potential neural injury, and the number of experimental animals needed ^95,96^. An inflammatory response was observed around the defect site at one month, likely due to the surgical adhesive. Cyanoacrylate, which is widely used for wound closure in clinical practice ^97^, can trigger an immune response ^97–99^. This assumption is supported by our six-month experiment, in which cyanoacrylate was not used, and no prominent inflammatory reaction was observed. These results also emphasize the polymers’ biocompatibility and suggest that the observed inflammation was primarily due to the adhesive rather than to the implanted materials. It is worth noting that NP bulging into the large created defect area was observed. True herniation, defined as NP displacement beyond the IVD boundary ^1^, did not occur, consistent with previous reports in sheep models ^92,100–102^. In contrast, human patients undergoing a discectomy procedure can experience symptomatic reherniation (18-27% of cases) ^103^. Although human and ovine IVDs are comparable, species differences in NP fluidity and fibrosity can explain the lower incidence of NP herniation in ovine IVDs ^92^. This inherent resilience to herniation may also explain the preservation of disc height at one month. Furthermore, only ∼80 mg of explanted tissue were removed in our study, whereas greater removals (100–200 mg of NP) have been associated with disc height loss in previous studies ^92^.

Partial or complete implant displacement was observed in vivo, reflecting the highly demanding mechanical environment of the disc. When fixation wings and PTFE membrane were used, both remained adherent to the adjacent vertebrae. However, due to their flexibility, the implants could still be pushed outward under intradiscal pressure. In the sutured group, the defect was closed by passing the thread twice through the outer AF layers without fully tightening the ends to avoid excessive tissue damage. Nevertheless, this approach was insufficient to prevent implant migration through the suture line. The addition of fixation wings and PTFE membrane appeared more effective than simple suturing, as portions of the implant remained within the AF defect, whereas in the latter case, the implant largely migrated outside the defect. Implant displacement remains a common challenge for AF repair materials, particularly when transitioning to large animal models ^61,104,105^. Despite this limitation, our implants supported cellular infiltration and neo-tissue formation, confirming their regenerative potential. Future studies should focus on improving implant stability, for instance, by using bio-adhesive fixation or suturing to the defect walls. Although polyester degradation produces acidic byproducts, these polymers have been previously used in IVD applications without reported cytotoxic effects ^106^. Still, further investigation into local pH changes and inflammatory responses during long-term degradation is needed.

Interestingly, the PCL and Blend implants were easily identifiable macroscopically after 6 months of in vivo implantation, with their lamellar structures preserved and clearly distinguishable. In contrast, the copolymer implants, when present, were primarily degraded and in some instances could not be distinguished macroscopically. These findings suggest that the copolymer underwent a predominantly bulk-erosion process, consistent with autocatalytic degradation occurring throughout the implant thickness. Such homogeneous erosion likely explains the rapid loss of multi-lamellar organization observed after six months. In contrast, the more crystalline PCL and blend materials exhibited slower, surface-driven degradation and retained their structural integrity. This observation correlates with our previous in vitro degradation study^107^: Although all polymers exhibited a decrease in molecular weight over time, only the copolymer showed visible nanofiber fractures and surface cracks under scanning electron microscopy. Furthermore, PCL and Blend polymer sheets maintained their tensile strength over six months, whereas the copolymer samples became brittle and their tensile force was unmeasurable at that time point. As we reported previously, the electrospun PCLA sheets displayed composition-dependent fiber diameters (ranging from 0.5 to 2.3 µm) and microstructural variations^107^ that may have influenced both in vivo degradation and cell infiltration. These morphological features, inherent to the copolymer versus blend architectures, likely contributed to the distinct degradation profiles observed between groups. In particular, the slower degradation of the blend, despite a higher PLA content, can be attributed to its biphasic morphology and higher crystallinity107, which limit water diffusion and hydrolytic chain scission, thereby delaying bulk erosion within the multilayer implant. Furthermore, the acidic and poorly buffered microenvironment of the intervertebral disc may have further accelerated the hydrolytic cleavage of ester bonds within the copolymer. In such confined conditions, the accumulation of acidic degradation products can promote autocatalytic hydrolysis, leading to a faster loss of molecular weight. This effect could be particularly pronounced for the homogeneous copolymer architecture, where degradation occurs throughout the bulk, whereas in more phase-segregated blends, degradation remains more surface-limited. Overall, this in vivo study is the first to demonstrate that PLA incorporation enhances PCL degradation within the IVD microenvironment, depending not only on PLA content but also on polymer structure (Blend vs. Copolymer).

A complete restoration of IVD mechanical function requires re-establishing NP hydration, for instance, through hydrogel injection. However, most current repair strategies focus solely on annulus fibrosus (AF) repair, leaving the nucleus pulposus (NP) untreated. In that case, the AF is exposed to higher mechanical stress, increasing the risk of implant failure and compromising long-term repair outcomes. Recent studies have highlighted the advantages of dual-target repair approaches that address both AF and NP, although such strategies remain in the early stages of development ^108–113^. These combined approaches have demonstrated superior outcomes compared with treatments targeting only one disc component. Nonetheless, hydrogel leakage remains a significant challenge, particularly when the AF defect is not adequately sealed.

Our degradation-tunable system could be further enhanced to synergistically release biological cues, such as anti-inflammatory ^114^, chemotactic, or growth factors ^89^.

## Conclusion

This study demonstrates the critical influence of PCLA polymer structure on its degradation behavior within the intervertebral disc microenvironment and its implications for annulus fibrosus repair. Aligned PCLA sheets supported in vitro AF cell alignment and proliferation. Ex vivo experiments confirmed the stable retention of the multi-lamellar implant within full-thickness annular defects and showed initial cell infiltration from native tissue. In vivo implantation in an ovine model showed good integration and tissue infiltration. Overall, the multi-lamellar, cell-free implant developed in this study successfully mimicked the native AF’s architecture and biomechanics, providing a supportive microenvironment for AF-like neo-tissue formation. PCL and blend implants retained their lamellar architecture for up to six months, whereas copolymers exhibited a more rapid yet controlled degradation. These findings indicate that tuning degradation kinetics through polymer design can provide an optimal balance between temporary mechanical support and progressive tissue regeneration. Further optimization of implant retention within the defect site is required to ensure long-term stability and reliable clinical translation.

## Acknowledgments

This study was supported by Fondation pour la Recherche Médicale (FRM, REP202110014192). The authors would like to thank the SC3M, Sc4Bio facilities from the Inserm/Nantes Université/ONIRIS UMR1229 RMeS Laboratory and SFR Bonamy, and the MicroPICell platforms from the University of Nantes and the IBISA MicroPICell facility (Biogenouest), member of the national infrastructure France-Bioimaging supported by the French national research agency (ANR-10-INBS-04). Furthermore, the authors wish to thank the personnel of the Preclinical Research and Investigation Center (CRIP) and the Medical Imaging Center at Oniris (O. Gauthier, P. Roy, P. Monmousseau, D. Rouleau, C. Raphael, S. Madec, A. Lafragette, and D. El-Hajj).

## Resource availability

### Lead contact

Requests for further information should be directed to and will be fulfilled by the lead contact, Catherine Le Visage.

### Materials availability

All reagents and protocols supporting the findings of this study are available from the lead contact upon request.

### Data and code availability

All data associated with this study are present in the paper or supplemental materials. Any additional information required to reanalyze the data reported in this paper will be available from the lead contact upon request.

### Author contributions

Conceptualization: CLV, JG, XG, H-J W, MF, CP, BN Methodology: MC, CP, CD, CF, CLV, JG, XG, CL, H-J W, MF Investigation: MC, CD, CF, PH, FE, MD, CL, A-K G-P, MV

Funding acquisition: CLV, JG, XG, H-J W, MF Project administration: CLV, XG, H-J W Supervision: CP, XG, CLV, H-J W, BN

Writing-original draft: MC, CLV

Writing-review & editing: MC, CLV, JG, CL

---

**Supplementary Figure 1.**
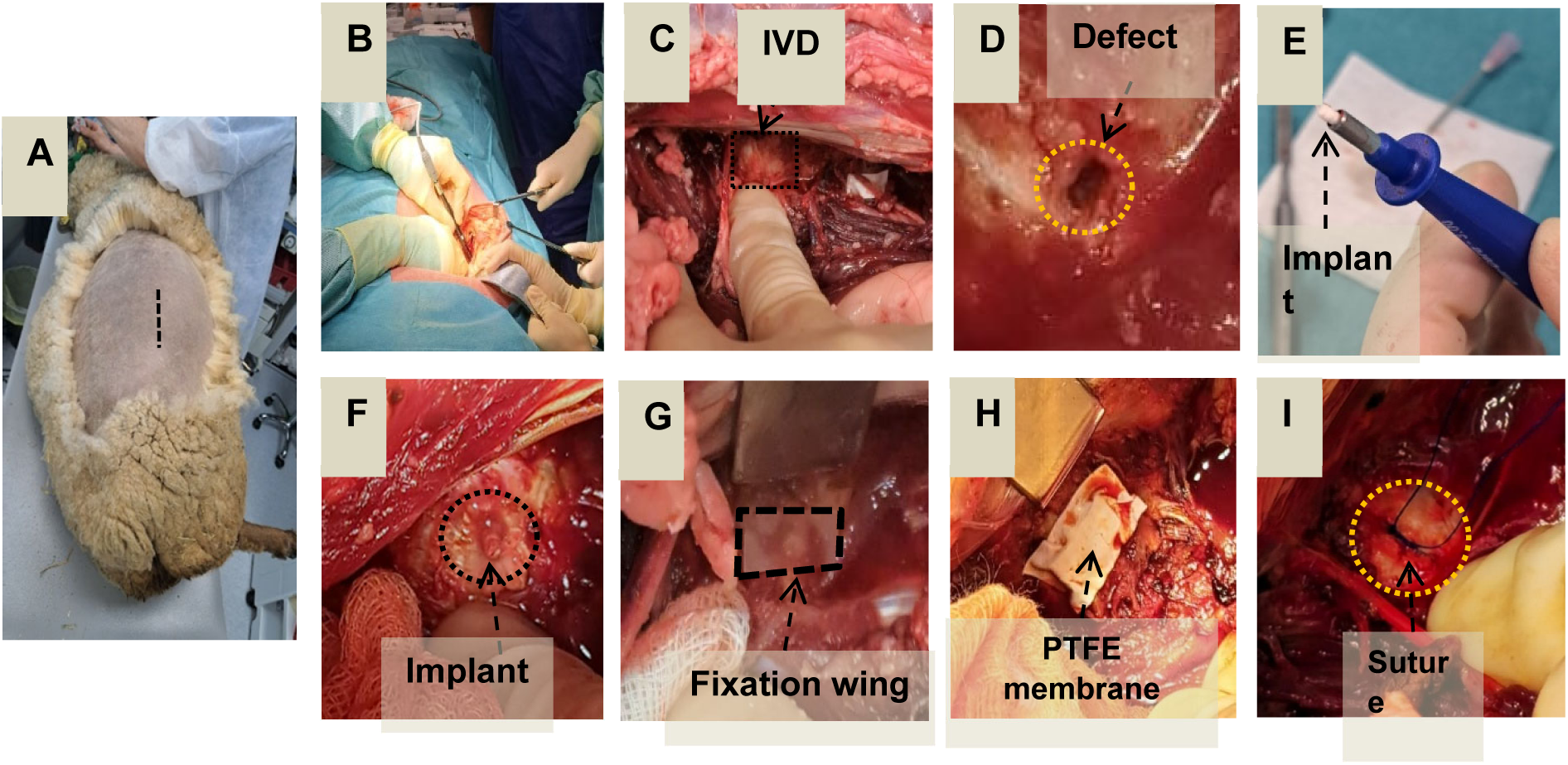
The implant can be placed within an annular defect through an anterolateral approach. The lumbar discs in an ovine model of simulated discectomy were used to evaluate implant repair efficacy in vivo at two time points: one month (n = 4 sheep) and six months. (n = 10 sheep). A total of five discs (L1–L2 to L5–L6) were used from each sheep. The discs were divided into the following experimental groups (n = 14 per group): an intact group, a non-implanted group, and three repaired groups based on the implant material (PCL, copolymer, or blend). (**A-I**) Illustration of the surgical procedure. (**A-C**) Accessing the disc via an anterolateral approach. (**D**) Defect creation using a 2 mm biopsy punch. (**E** and **F**) Placement of the implant guided with a 3 mm biopsy punch. (**G** and **H**) Gluing the fixation wings (**G**) and a PTFE patch (**H**) to the adjacent vertebrae. (**I**) Suturing the defect for the six-month time point.

**Supplementary Figure 2.**
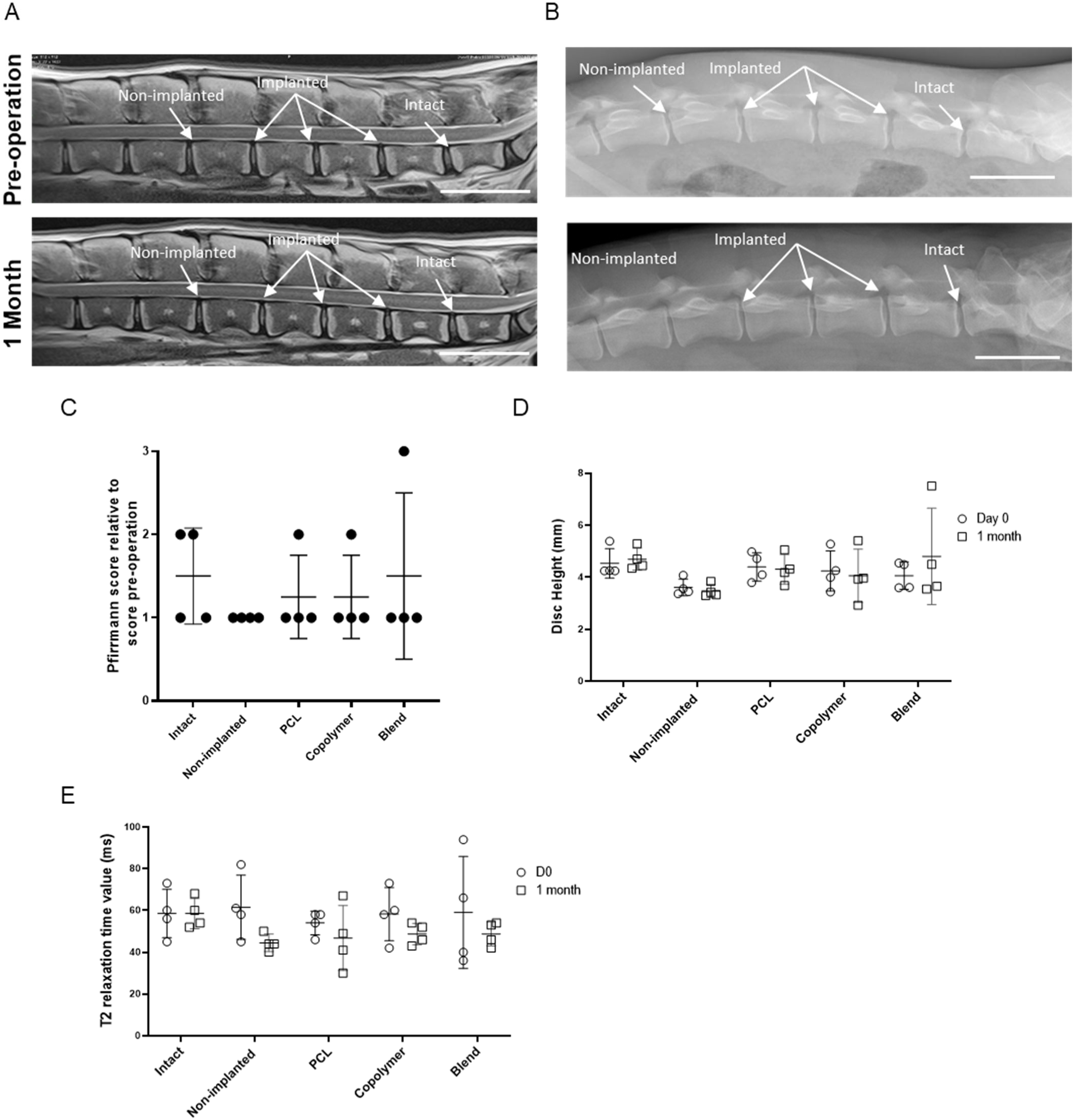
MRI and X-ray imaging results of the ovine discs at 1 month showed no signs of degeneration. MRI and X-ray scans were performed preoperatively (Day 0) and pre-euthanasia (1 month). Five discs (L1–L2 to L5–L6) were used from each sheep (n = 4). The discs were divided into the following experimental groups (n=4 per group): an intact group, a non-implanted group, and three repaired groups based on the implant material (PCL, copolymer, or blend). The images were analyzed using OsiriX software®. (**A** and **B**) Representative sagittal T2 MRI (**A**), and X-ray images (**B)** of the lumbar discs. Scale bar: 50 mm (**C**) Pfirrmann grading of the discs. (**D**) Disc height values. (**E** T2 relaxation time values. All data are represented as mean ± standard deviation . N = 4 discs per group. Comparisons among groups were performed using one way (Pfirrmann score) or two-way ANOVA (Disc height, T1, T2 and T2*), followed by a Tukey’s test. ∗ = statistically significant; p ≤ 0.05. PCL: polycaprolactone, Copolymer: C-CL_90%_LA_10%_, Blend: B-PCL_80%_PLA_20%_, MRI: magnetic resonance imaging.

**Supplementary Figure 3.**
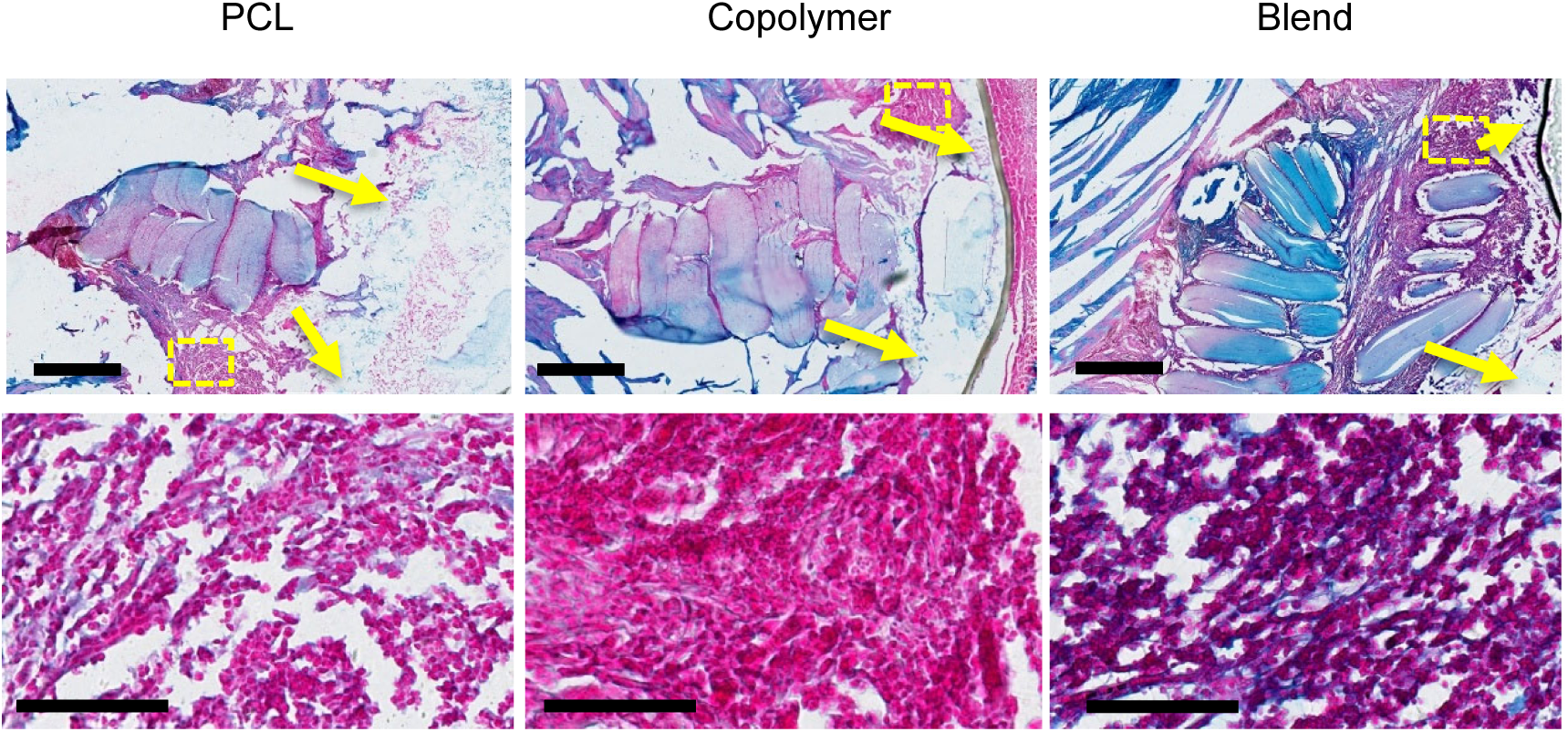
Inflammatory tissue around the implant. Representative images of Alcian Blue-stained sections of the discs are shown. Yellow arrows indicate the location of the glue material. Dotted lines indicate the location of the high magnification image. Scale bar: 1 mm for the low magnification images (top row) and 100 μm for high magnification images (bottom row).

**Supplementary Figure 4.**
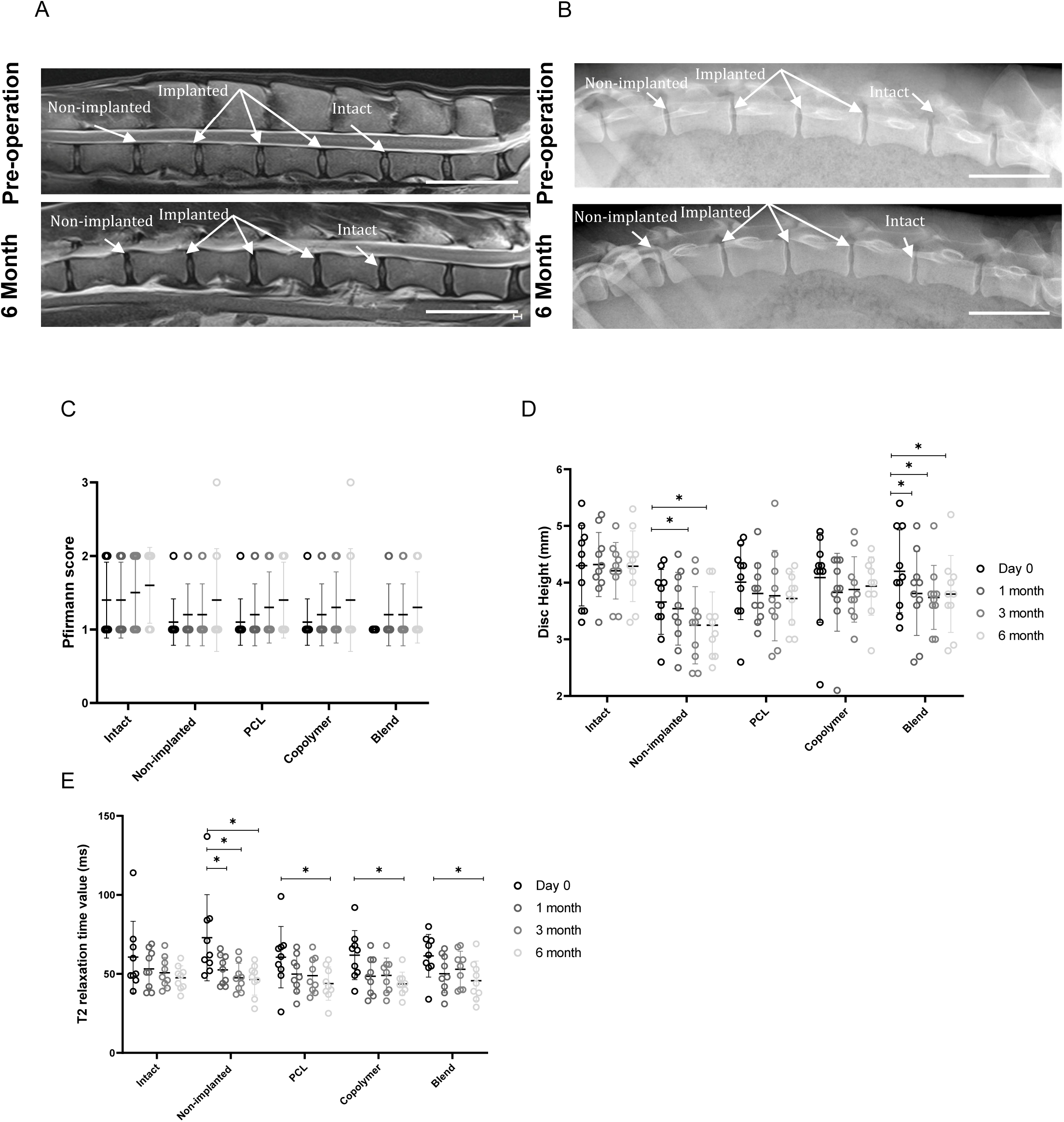
MRI and X-ray imaging results of the ovine discs at 6 month showing delayed disc dehydration for the repaired groups. MRI and X-ray scans were performed preoperatively (Day 0) and at one, three and six. Five discs (L1–L2 to L5–L6) were used from each sheep (n = 10). The discs were divided into the following experimental groups (n = 4 per group): an intact group, a non-implanted group, and three repaired groups based on the implant material (PCL, copolymer, or blend). (**A** and B) Representative sagittal T2 MRI (**A**) and X-ray images (**B)** of the lumbar discs. Scale bar: 50 mm (**C**) Pfirrmann grading of the discs. (**D**) Disc height values. (**E**) T2 relaxation time values. All data are represented as mean ± standard deviation . N = 10 discs per group. Comparisons among groups were performed using two-way ANOVA, followed by a Tukey’s test. ∗ = statistically significant; p ≤ 0.05. PCL: polycaprolactone, Copolymer: C-CL_90%_LA_10%_, Blend: B-PCL_80%_PLA_20%_, MRI: magnetic resonance imaging.

**Supplementary Figure 5.**
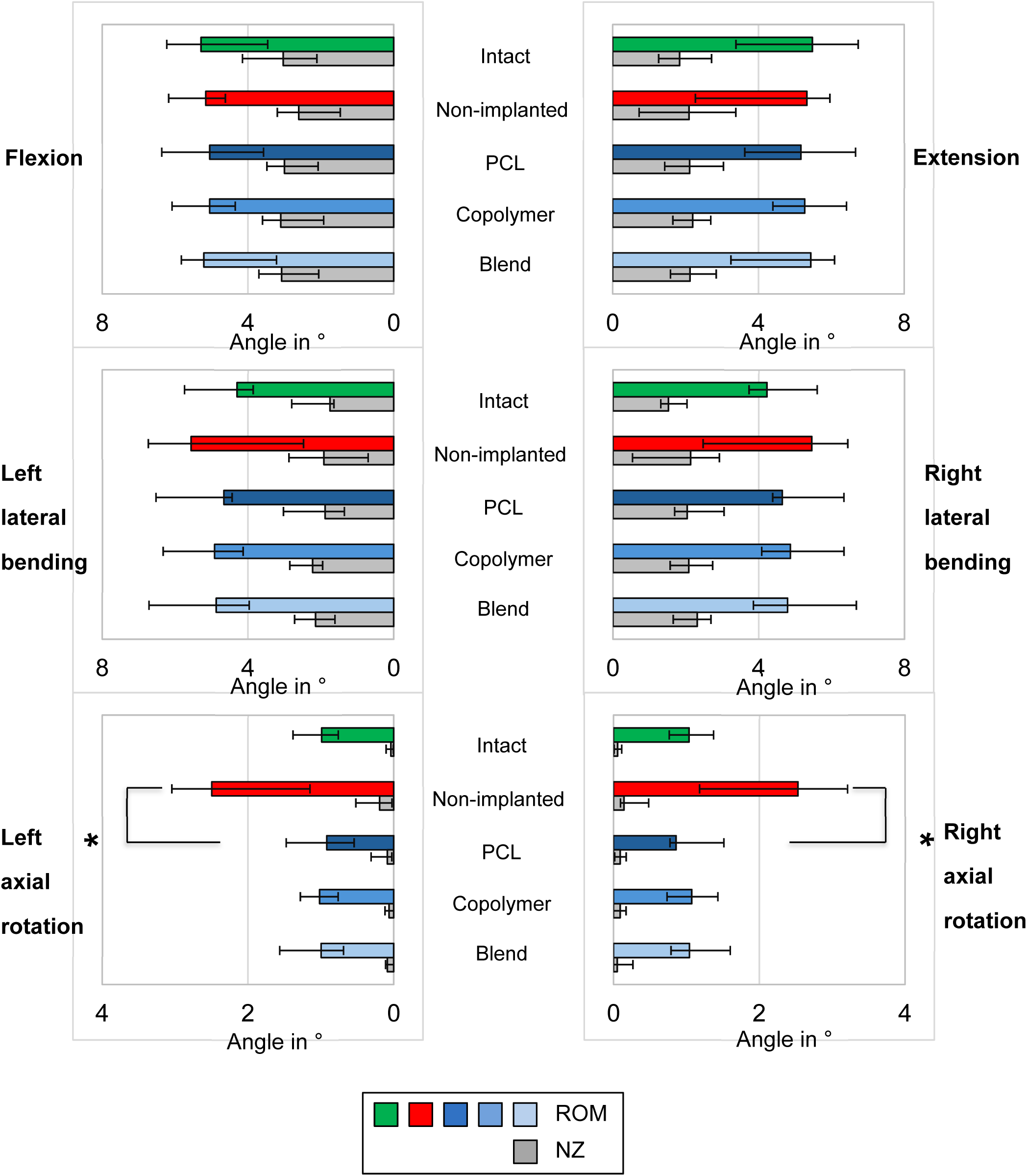
Ex vivo biomechanical testing of the ovine discs at 6 month. Five discs (L1–L2 to L5–L6) were analyzed from each sheep (n = 6). The discs were divided into the following experimental groups (n = 6 per group): an intact group, a non-implanted group, and three repaired groups based on the implant material (PCL, copolymer, or blend). The specimens were loaded with displacement-controlled (1 °/s) pure moments of 7.5 Nm 73 in flexion/extension, lateral bending, and axial rotation using a well-established spine tester to determine range of motion (ROM) and neutral zone (NZ). All data are represented as mean ± standard deviation . N = 6 discs per group. Comparisons among groups were performed using pairwise Kruskal-Wallis test with Dunn-Bonferroni post-hoc correction for multiple groups with low sample size. ∗ = statistically significant; p ≤ 0.05. PCL: polycaprolactone, Copolymer: C-CL_90%_LA_10%_, Blend: B-PCL_80%_PLA_20%_, ROM: range of motion, NZ: neutral zone.

**Supplementary table 1.** AF cells alignment.

| Condition | Alignment score | Dispersion |
| --- | --- | --- |
| 2D | $0.46 \pm 0.1$ | $23.9 \pm 8.2^\circ$ |
| PCL | $0.97 \pm 0.0$ | $15.7 \pm 1.2^\circ$ |
| Copolymer | $0.96 \pm 0.0$ | $15.5 \pm 1.8^\circ$ |
| Blend | $0.97 \pm 0.0$ | $18.8 \pm 3.2^\circ$ |

\* The dispersion ( $^\circ$ ) describes the standard deviation of the Gaussian, and the alignment score describes the goodness of the alignment. from 0: no preferred orientation to 1: aligned. (N = 1, n = 3)

**Supplementary table 2.** Summary of in vivo outcome at one month.

| Polymer type | Located within or around the defect | Extruded Outside the defect | Sign of degradation |
| --- | --- | --- | --- |
| PCL | 4/4 | 4/4 | 0/4 |
| Copolymer | 3/4 | 3/3 | 0/3 |
| Blend | 3/4 | 3/3 | 0/3 |

**Supplementary table 3.** Summary of in vivo outcome at six month.

| Polymer type | Located within or around the defect | Extruded Outside the defect | Sign of degradation |
| --- | --- | --- | --- |
| PCL | 9/10 | 10/10 | 0/10 |
| Copolymer | 4/10 | 4/4 | 4/4 |
| Blend | 9/10 | 10/10 | 0/10 |

